# Morphological Variation and DUS Characterization in Bitter Gourd (*Momordica charantia* L.) collections from Eastern India for Varietal Identification and Genetic Improvement

**DOI:** 10.1101/2025.02.13.638034

**Authors:** Anita Verma, Alpana Joshi

## Abstract

This study investigates the morphological variation and Distinctness, Uniformity, and Stability (DUS) characterization of *Momordica charantia* collections from Eastern India to facilitate varietal identification and genetic improvement. Twenty-two Bitter Gourd varieties were characterized based on morphological traits to assess genetic diversity and enhance breeding program efficiency. Significant variations were observed in fruit morphology (colour, length, surface tubercles, and ridges) and seed characteristics (Colour and margin indentation) enabling effective varietal differentiation. Notably, four accessions were classified as *M. charantia var. muricata* due to their distinctly small fruit size, highlighting the presence of diverse morphological forms within the studied population. Additionally, admixture was detected in seven varieties, which underscores the necessity for purification processes to ensure the commercial viability of these accessions. This research emphasizes the rich genetic diversity of Bitter Gourd in Eastern India and identifies key traits that are essential for distinguishing between varieties. The findings provide valuable insights for breeders to develop improved cultivars with enhanced productivity and consumer-preferred traits. Moreover, this study supports ongoing plant breeding efforts to enhance crop quality and yield by identifying reliable morphological traits for varietal differentiation. Furthermore, it recommends additional research to refine DUS criteria by incorporating more traits that effectively differentiate between smooth-type Bitter Gourd varieties.

## INTRODUCTION

Bitter Gourd (*Momordica charantia* L.), also known as karela or bitter melon, is a diploid species (2n = 22) with an estimated genome size of 339 Mb (Urasaki *et al*., 2017) It is a valuable member of the Cucurbitaceae family cultivated extensively across tropical and subtropical regions of Asia, South America, East Africa, and the Caribbean. Bitter Gourd is renowned for its dual significance as a vegetable and medicinal plant (Cui *et al*., 2020). The exact origin of Bitter Gourd (*Momordica charantia*) has been debated for long. While earlier studies indicated that Eastern Asia might be the centre of origin, recent advancements in molecular research, such as those conducted by Schaefer and Renner (2010), have proposed that Africa is the primary birthplace of the entire *Momordica* genus. It is believed that Asian species, including Bitter Gourd, originated from a single long-distance dispersal event from Africa approximately 19 million years ago. This suggests that monoecious *Momordica* species, like Bitter Gourd, likely evolved from their dioecious ancestors’ native to Africa.

Despite its African origin, India is widely considered a likely centre of diversity for Bitter Gourd. This hypothesis is supported by early written records from India, the coexistence of domesticated and wild progenitors within the country, and its early references in Ayurveda literature (Sivarajan & Balachandran, 1994; Decker-Walters, 1999). Although China has historical records of Bitter Gourd cultivation, these records appear later than those in India, suggesting a possible introduction from India (Yang & Walters, 1992). Irrespective of its precise centre of domestication, this crop is renowned across regions in Asia for its numerous health benefits. It boasts a long history of use in traditional medicine, with applications for ulcers, skin conditions, and diabetes (Nesar Ahmad *et al*., 2016). Recent research has significantly strengthened the recognition of Bitter Gourd’s health benefits, demonstrating its potential in lowering blood sugar levels in type 2 diabetes, exhibiting antiviral and antibiotic properties, and offering overall health benefits (Jia *et al*., 2017). The increasing recognition of Bitter Gourd as a natural and health-promoting food option has driven a surge in consumer demand, spurring significant research and development efforts. As a result, breeding programs are focusing on developing improved hybrids that address market demands for higher yields and superior fruit quality while meeting consumer preferences for enhanced nutritional and medicinal benefits.

The foundation of a successful breeding program lies in the efficient utilization of existing genetic diversity. Breeders require access to diverse plant genetic resources, supported by detailed and reliable information, to advance their work. Distinctiveness, Uniformity, and Stability (DUS) testing, as mandated by the Protection of Plant Varieties and Farmers’ Rights (PPVFR) Act 2001, plays a crucial role in modern plant breeding through several key functions. Firstly, DUS testing serves as the foundation for protecting intellectual property rights through Plant Variety Protection (PVP) systems, safeguarding the interests of breeders and encouraging innovation. Secondly, it ensures the integrity of new cultivars by rigorously assessing their unique traits, consistent performance, and stability over generations. Thus, precise genotype characterization is essential. DUS testing determines whether a newly developed cultivar is distinct from existing ones within the same species and prevents redundancy among gene bank accessions (Mahapatra *et al*., 2022; Mallikarjuna *et al*., 2023).

Moreover, information on the available genetic variability for different quantitative traits is invaluable for devising effective breeding strategies for genetic enhancement. Continuous efforts in this direction will be crucial for developing cultivars with improved productivity and consumer preference. Despite its importance, research on the characterization of available Bitter Gourd (*Momordica charantia* L.) genotypes remains limited. Given the critical need for this information in both genetic resource management and breeding programs, this study aimed to characterize and catalogue the unique traits of 22 landraces collected primarily from Odisha and West Bengal, regions within eastern India — a recognized centre of diversity for Bitter Gourd. Understanding the vast variability within this gene pool is vital for breeding advancements. To ensure comprehensive characterization and seamless data exchange, the study employed standardized descriptor lists provided by Bioversity International (2007). This well-established system facilitates the efficient utilization of these valuable genetic resources by providing a common framework for data collection and sharing among researchers and breeders worldwide.

## MATERIALS AND METHODS

### Experimental Site and Design

This study was conducted during the Kharif season of 2020 at the BRDC Centre of Bayer Crop Science, located in Kallinayakanhalli, Bangalore, and Karnataka, India. Twenty-two diverse Bitter Gourd genotypes (Table 1) were evaluated in a randomized complete block design (RCBD) with three replications, with each replication comprising five plants per genotype.

**Table 1.** List of Varieties used for DUS testing.

| **Code** | **Land race name** | **Code** | **Land race name** |
| --- | --- | --- | --- |
| BG-001 | Kajal | BG-012 | Chaman |
| BG-002 | Kuheli | BG-013 | Bolder Uchaa |
| BG-003 | Kalisankar | BG-014 | Diamond |
| BG-004 | RS Meghana | BG-015 | 2015 Max |
| BG-005 | Tiyasha | BG-016 | Kajalgouri |
| BG-006 | Kalasona | BG-017 | SP Dhanakali |
| BG-007 | Nilima | BG-018 | BG-OP-002 |
| BG-008 | BG-OP-008 | BG-020 | Bolder |
| BG-009 | Katahi | BG-021 | Nilisha Gold |
| BG-010 | SP Daspur | BG-022 | Tiyasha Double |
| BG-011 | MJ5 Jonpuri | BG-023 | MJ5 Summer special |

### Seedling Preparation and Transplantation

Seedlings were raised in a controlled environment within a hi-tech greenhouse nursery. Seeds were sown in plastic pro-trays filled with a sterile, soilless medium comprising a 4:1 mixture of coco peat and vermicompost. This substrate provided a uniform and consistent environment for germination and seedling establishment. Prior to sowing, seeds were treated with a fungicide solution to minimize the risk of damping-off diseases and ensure healthy seedling development. Twenty-day-old seedlings were carefully transplanted into the experimental field, maintaining a spacing of 45 cm between plants and 1.5 meters between rows. Standard agronomic practices, including irrigation, fertilization, and weed management, were consistently applied throughout the crop cycle to ensure optimal plant growth and development.

### Data Collection

To assess the genetic diversity and characterize the genotypes, 29 morphological traits (Table 2) were systematically recorded for each genotype across various growth stages. These observations were conducted in accordance with the DUS test guidelines outlined by the Protection of Plant Varieties and Farmers’ Rights Authority (PPV and FR Act., 2001). Key morphological traits assessed during the DUS test included fruit, vegetative, flower, and seed characteristics. Fruit traits included length, diameter, and skin colour at maturity, and shape, such as oblong or spindle. Vegetative traits included vine length, stem shape, internode length, leaf blade length, margin type, and the number of lobes. Flower traits were evaluated for colour and ovary length on the day of anthesis. Seed traits comprised the number of seeds per fruit, along with seed length, colour, and surface texture.

**Table 2.** List of Traits Assessed as per DUS Guidelines.

| S.N. | Trait | S.N. | Trait |
| --- | --- | --- | --- |
| 1 | Cotyledon: intensity of green<br>Colour | 16 | Peduncle: length |
| 2 | Plant: main vine length | 17 | Fruit: colour of skin |
| 3 | Stem: shape | 18 | Fruit: shape of base at<br>peduncle end |
| 4 | Stem: length of internodes of main<br>stem (between 15<sup>th</sup> - 20<sup>th</sup> node) | 19 | Fruit: shape of apex at<br>blossom end |
| 5 | Stem: number of primary branches | 20 | Fruit: shape in longitudinal<br>section |
| 6 | Leaf blade: length | 21 | Fruit: tubercles |
| 7 | Leaf blade: margin | 22 | Fruit: tubercles prominence |
| 8 | Leaf blade: shape | 23 | Fruit: ridge |
| 9 | Leaf blade: number of lobes | 24 | Fruit: Colour of skin at ripe<br>stage |
| 10 | Leaf blade: depth of lobbing | 25 | Seediness (no. of<br>seeds/fruit) |
| 11 | Petiole: length | 26 | Seed: length |
| 12 | Flower colour | 27 | Seed: colour |
| 13 | Ovary: length<br>(on the day of anthesis) | 28 | Seed: indentation of margin |
| 14 | Fruit: length | 29 | Seed surface |
| 15 | Fruit: diameter |  |  |

## RESULTS

In the present investigation, 29 traits were assessed across various plant aspects, including plant characteristics, leaf morphology, fruit characteristics, and seed morphology (Table 3 and 4). Among these traits, seven were identified as invariant across all 22 genotypes examined. The consistent traits included the length of the main vine, which was uniformly short and viny, measuring less than 2 meters; stem shape, consistently observed as angular; and leaf blade shape, which was cordate with five distinct and deeply lobed segments. Additionally, the number and depth of lobes exhibited no genotypic variation, and flower colour was uniformly yellow across all genotypes. Seed surface texture was also conserved, being rough in all samples. In contrast, the remaining traits displayed significant variation among the genotypes, reflecting the inherent genetic diversity within the population. These variable traits have been further categorized based on their biological relevance into three primary domains: plant characteristics, which include growth habit, branching pattern, and other structural traits; fruit characteristics, encompassing attributes such as size, shape, and colour; and seed characteristics, including metrics related to size, weight, colour, and surface patterns. This categorization facilitates a focused discussion of the observed phenotypic diversity and its potential implications for genotype characterization and breeding programs.

**Table 3.**
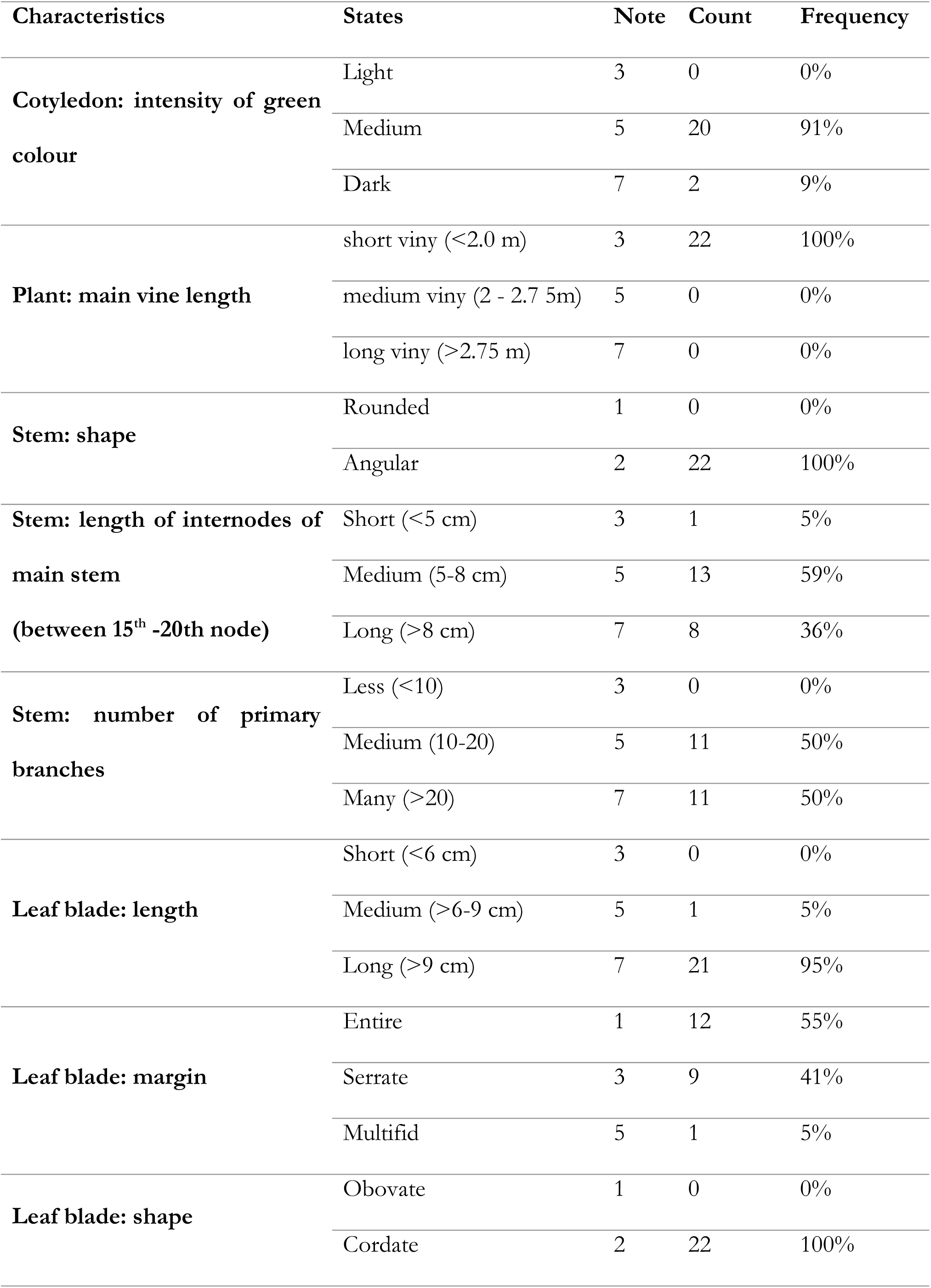
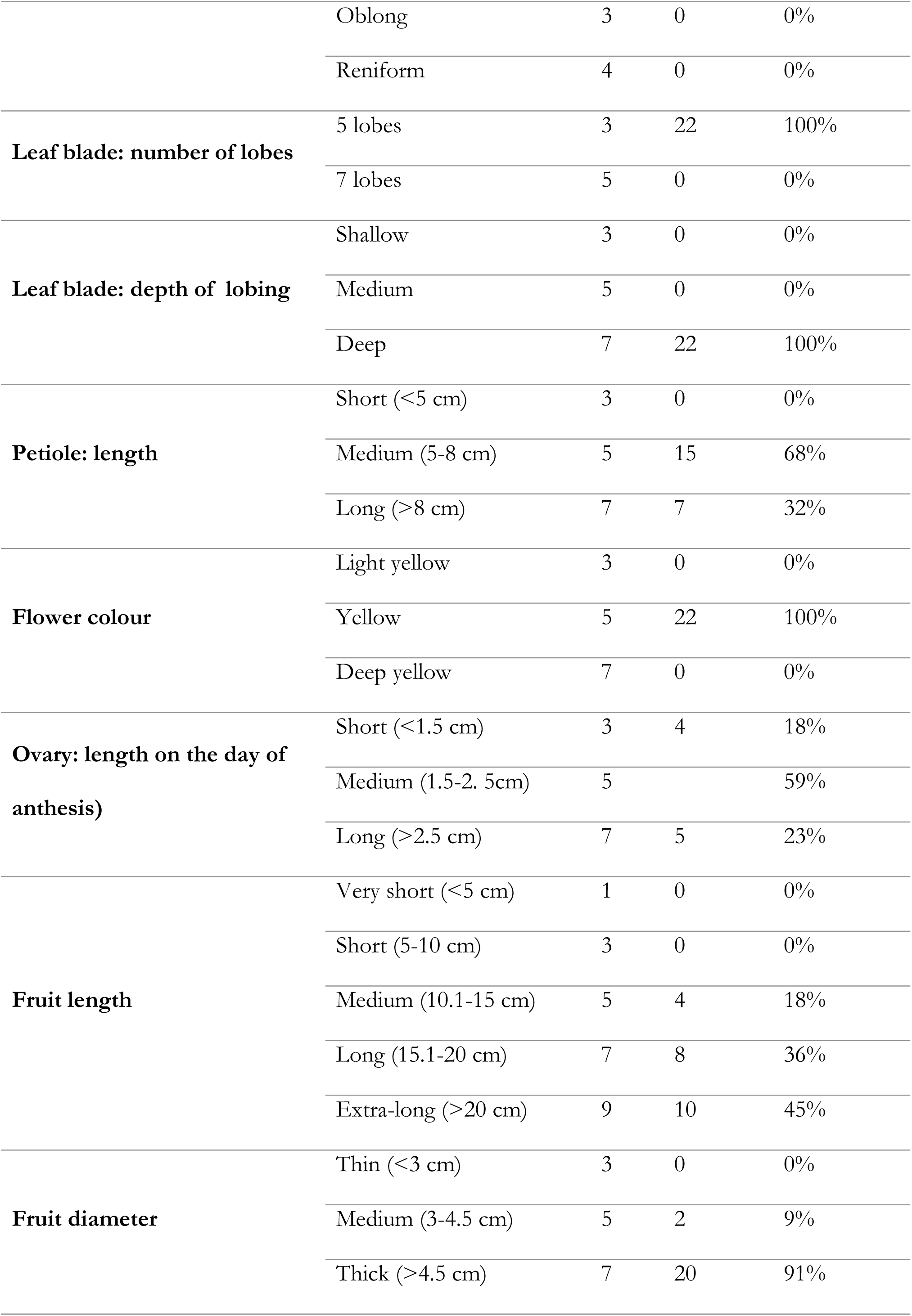
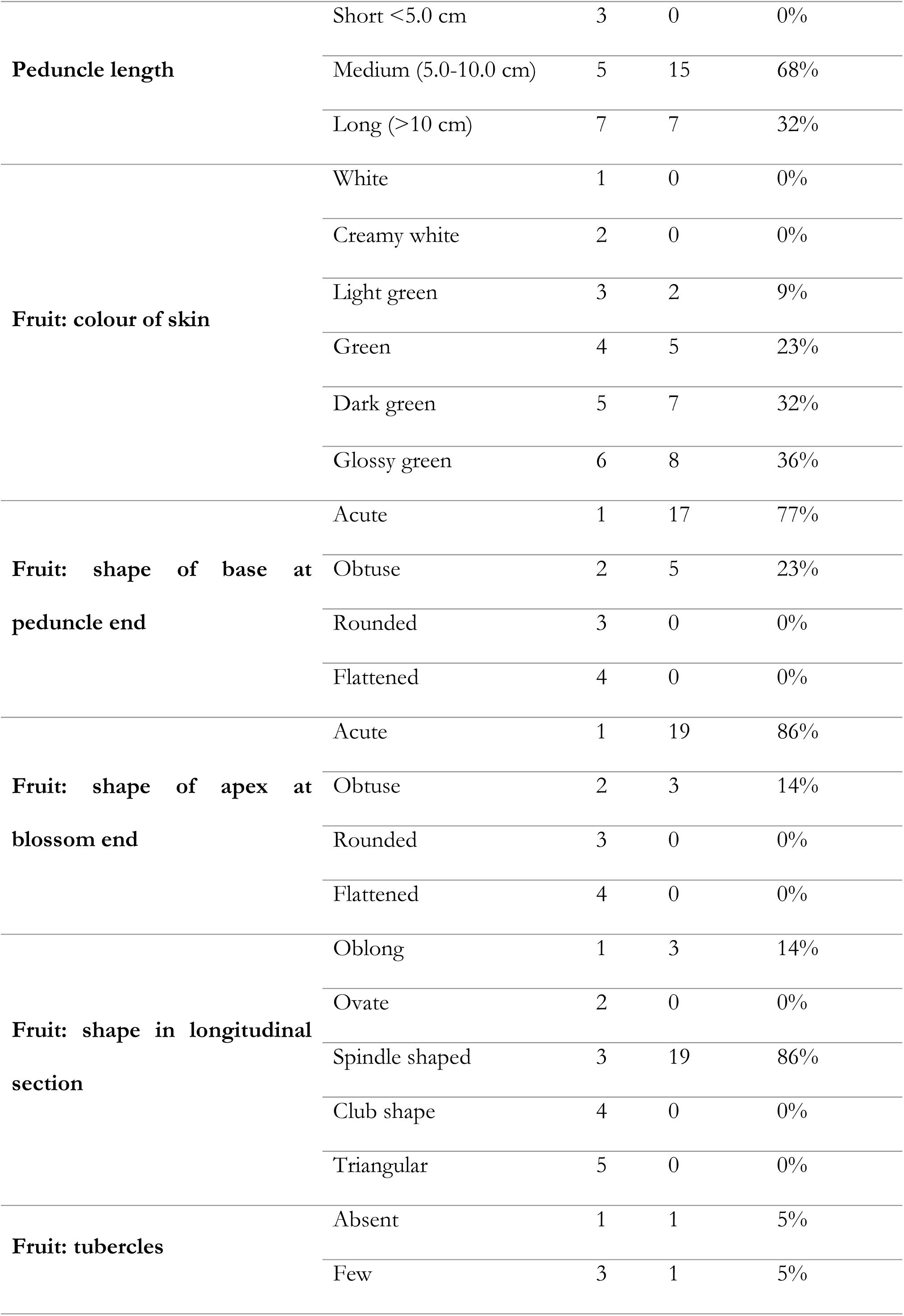
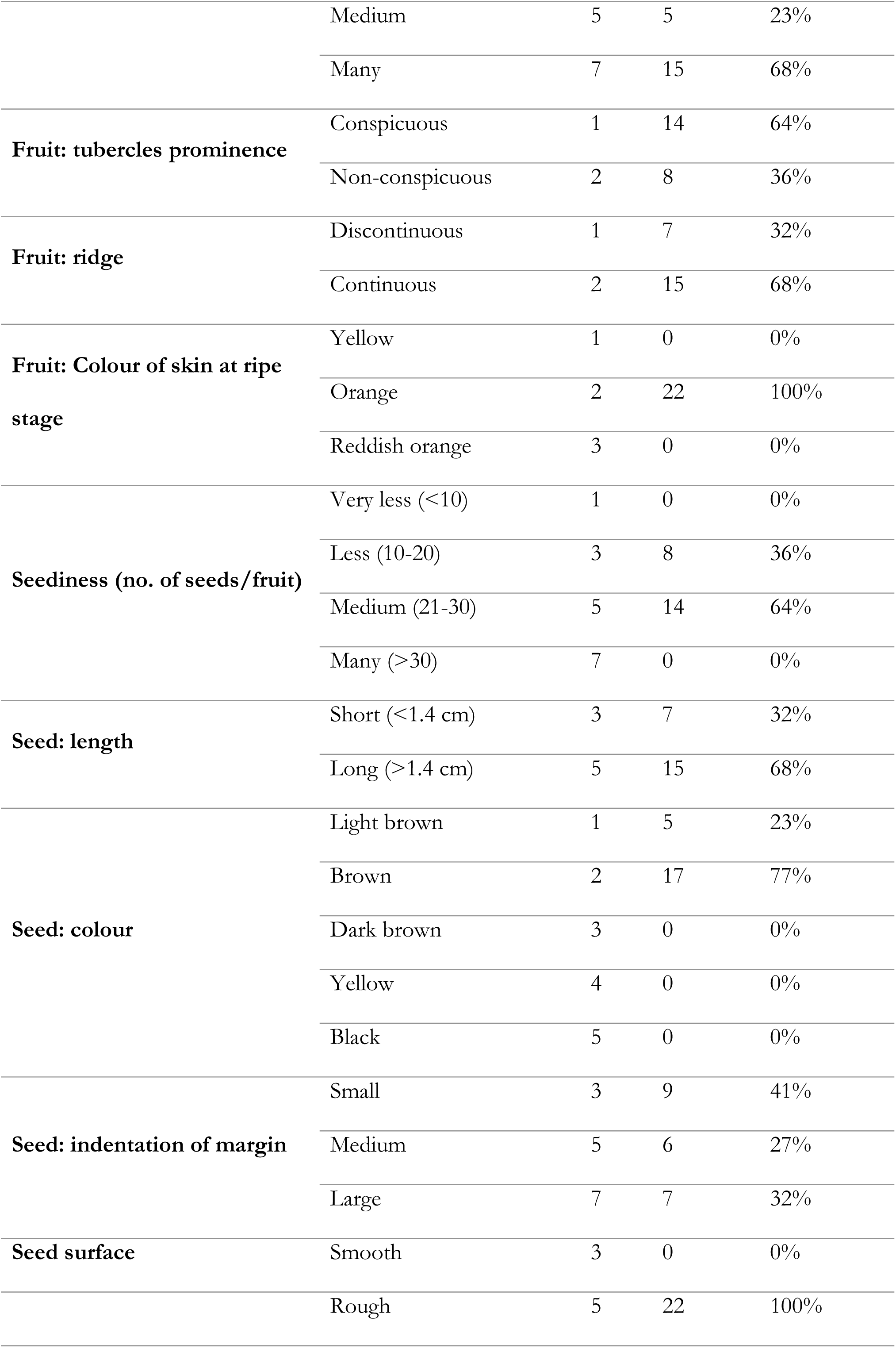
Frequency of observed variation for each trait based on DUS Guideline.

**Table 4.** Characterization of Bitter Gourd Genotypes for Plant Characteristics based on traits as per the DUS Guideline.

| **Genotypes** | **Cotyledon:**<br>**intensity**<br>**of green**<br>**Colour** | **Stem: length**<br>**of**<br>**internodes**<br>**of main**<br>**stem** | **Stem:**<br>**number**<br>**of**<br>**primary**<br>**branches** | **Leaf**<br>**blade:**<br>**length** | **Leaf**<br>**blade:**<br>**margin** | **Petiole:**<br>**length** | **Ovary:**<br>**length** |
| --- | --- | --- | --- | --- | --- | --- | --- |
| Kajal | Medium | Medium | Medium | Long | Serrate | Medium | Medium |
| Kuheli | Medium | Long | Medium | Long | Entire | Long | Long |
| Nilima | Dark | Medium | Medium | Long | Entire | Medium | Medium |
| Tiyasha | Medium | Medium | Medium | Long | Entire | Medium | Long |
| BG-OP-008 | Medium | Medium | Medium | Medium | Entire | Medium | Long |
| BG-OP-002 | Medium | Long | Medium | Long | Serrate | Medium | Long |
| Katahi | Medium | Long | Medium | Long | Entire | Medium | Medium |
| Bolder Uchaa | Medium | Medium | Medium | Long | Entire | Medium | Medium |
| Kalasona | Medium | Short | Medium | Long | Entire | Medium | Medium |
| RS Meghana | Medium | Medium | Medium | Long | Entire | Medium | Short |
| 2015 Max | Medium | Medium | Many | Long | Serrate | Long | Medium |
| Bolder | Medium | Medium | Many | Long | Entire | Medium | Short |
| Diamond | Medium | Medium | Many | Long | Entire | Medium | Short |
| Kajalgouri | Dark | Medium | Many | Long | Serrate | Medium | Medium |
| Kalisankar | Medium | Medium | Many | Long | Serrate | Medium | Medium |
| Nilisha Gold | Medium | Long | Medium | Long | Serrate | Medium | Medium |
| SP Daspur | Medium | Medium | Many | Long | Entire | Long | Medium |
| SP Dhanakali | Medium | Medium | Many | Long | Serrate | Long | Medium |
| Tiyasha<br>Double | Medium | Long | Many | Long | Serrate | Medium | Long |
| Summer<br>special | Medium | Long | Many | Long | Multifid | Long | Short |
| Chaman | Medium | Long | Many | Long | Entire | Long | Medium |
| Jonpuri | Medium | Long | Many | Long | Serrate | Long | Medium |

### Plant Characteristics

Cotyledon colour was assessed across the collected germplasm, revealing that 91% of the evaluated material exhibited a medium cotyledon Colour, whereas 9% (2 out of 22 genotypes) displayed a darker pigmentation (Table 3 and 4). This trait, though not directly linked to yield, provides a morphological marker that can facilitate genotype differentiation. Key yield-contributing traits such as vine length, the number of productive branches, and inter-nodal distance between fruiting nodes were also evaluated. These traits significantly influence yield potential by contributing to an increased number of branches, flowers, and fruits. The vine length for all genotypes under study was uniformly small, measuring less than 2 meters, indicating compact growth habit suitable cultivation systems. However, substantial variation was noted in the number of productive branches and inter-nodal distance among the genotypes. The inter-node length of the main stem, measured between the 15th and 20th nodes, ranged from short (<5 cm) to long (>8 cm). Among the genotypes, Kalasona exhibited a unique short inter-nodal distance of less than 4 cm, whereas all other genotypes had either medium (5–8 cm) or long (>8 cm) inter-nodal distances. This variability in inter-nodal length could provide potential selection criteria for breeding programs aiming to optimize plant architecture for higher yields. The distribution of primary branches across the collection demonstrated a balanced frequency, with 50% of the genotypes categorized as medium (10–20 branches) and the remaining 50% classified as having many branches (>20 branches) (Table 3). Notably, none of the genotypes exhibited fewer than 10 branches, highlighting their potential as productive cultivars or promising material for breeding programs targeting increased yield efficiency. Leaf traits, while not directly contributing to yield, play an integral role in genotype identification and differentiation. Leaf blade length was predominantly long (>9 cm), observed in 95% of the genotypes. Only one genotype (5%) displayed a medium leaf blade length (6–9 cm). Significant variability was recorded in leaf blade margin types, ranging from entire to serrate and multifidi. While 95% of the genotypes exhibited entire or serrate margins, the genotype Summer Special uniquely displayed a multifid leaf margin, providing a distinctive morphological feature. Petiole length, another crucial leaf trait, was predominantly medium (5–8 cm) in 68% of the genotypes, whereas 32% (7 out of 22 genotypes) exhibited long petioles (>8 cm). This diversity in petiole length, along with other leaf morphological traits, underscores the phenotypic variation present within the collection and offers a basis for selecting desirable traits in breeding programs (Tables 3 and 4). This analysis highlights the genetic diversity and phenotypic variability in key agronomic and morphological traits, suggesting that the evaluated germplasm possesses potential for cultivar improvement and the development of high-yielding varieties.

### Fruit Characteristics

A notable variation in fruit length was found among 22 Bitter Gourd collections, ranging from 11 cm to 25 cm. Of these, 36% (8 collections) measured between 15 cm and 20 cm, while 45% were extra-long, at 20 cm to 25 cm. Collections like Bolder Uccha, Diamond, Bolder, and 2015 Max had lengths of 10 cm to 12 cm, classifying them as medium-length per DUS guidelines. Regarding fruit Colour, all possible Colour variations were observed, ranging from light green to glossy green (Table 5; Figure 1). Among the 22 varieties, Kuheli and Kalasona were the only ones that displayed a white fruit Colour. Five varieties were classified as green, seven were dark green, and eight exhibited a glossy green appearance. This broad spectrum of fruit shape and colour highlights the significant genetic variability present within Bitter Gourd, offering substantial opportunities for seed companies to further enhance desirable traits. Based on the combination of fruit length and colour, the 22 collections were categorized into 10 distinct groups (Table 5). Given the increasing market preference for dark green and glossy green Bitter Gourd, the presence of such variations, along with diverse length options, is advantageous for breeders aiming to develop improved cultivars

**Table 5.** Classification of Genotypes groups based on the Colour and Length.

| S.N. | Length and Colour Group | Genotype(s) |
| --- | --- | --- |
| 1 | Extra-long Dark Green | BG-OP-008, Kajal Gouri, SP Dhanakali, Jonpuri |
| 2 | Extra-long Glossy Green | BG-OP-002, Nilisha Gold, Tiyasha Double, Chaman |
| 3 | Extra-long Green | Katahi |
| 4 | Extra-long Light Green | Kuheli |
| 5 | Long Dark Green | Kajal, Kalisankar, SP Daspur |
| 6 | Long Glossy Green | Summer special |
| 7 | Long-Green | Nilima, Tiyasha, RS Meghana |
| 8 | Long-Light Green | Kalasona |
| 9 | Medium-Glossy Green | Bolder Uchaa, 2015 Max, Bolder |
| 10 | Medium-Green | Diamond |

**Figure 1.**
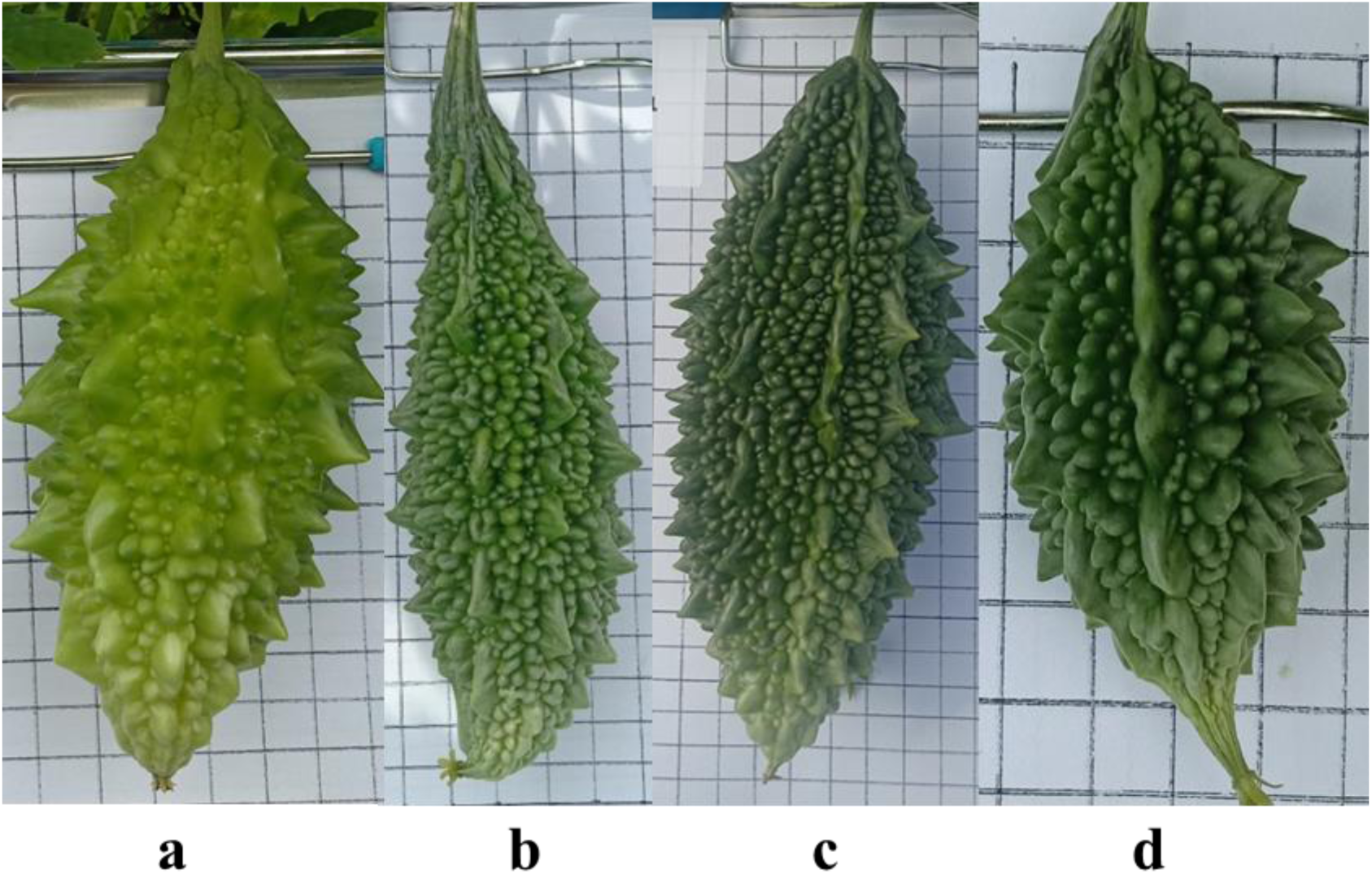
Variation in Fruit Colour: (a) Light Green, (b) Green, (c) Dark Green, (d) Glossy

Notably, only four genotypes were classified as medium-length, with fruit sizes ranging between 10 cm and 12 cm (Figure 2). These four types could be identified as belonging to *Momordica charantia var. muricata*, not only based on their length but also considering additional morphological characteristics such as fruit girth and shape. In terms of fruit diameter, only two varieties—Bolder and Diamond—exhibited a diameter between 3 cm and 4.5 cm. The remaining 20 varieties fell into the thick category, with fruit diameters exceeding 4.5 cm (Table 6).

**Figure 2.**
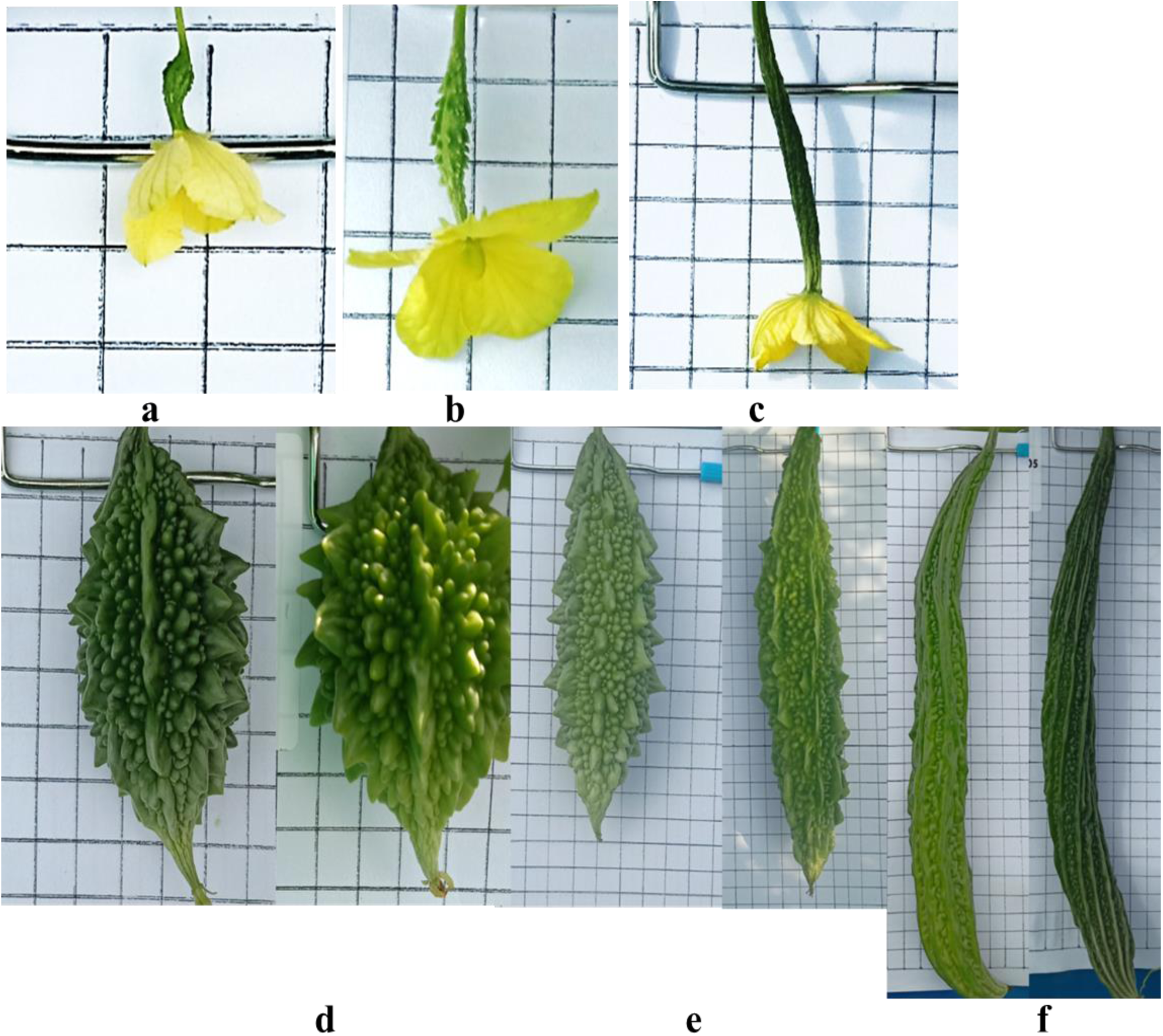
Variation in Ovary and Fruit Length: (a, d) Short, (b, e) Medium, and (c, f) Long.

**Table 6.** Characterization of Bitter gourd Genotypes for Fruit Characteristics based on traits as per the DUS Guideline.

| Characteristics | Fruit length | Peduncle length (cm) | Fruit: colour of skin | Fruit: shape of base at peduncle end | Fruit: shape of apex at blossom end | Fruit: tubercles prominence | Fruit: shape in longitudinal section | Fruit: tubercles | Fruit: ridge | Fruit: colour of skin at ripe stage |
| --- | --- | --- | --- | --- | --- | --- | --- | --- | --- | --- |
| Kajal | long | 5-10 | Dark Green | Acute | Acute | Conspicuous | Oblong | Medium | Continuous | Orange |
| Kuheli | Extra long | >10 | Light Green | Obtuse | Acute | Conspicuous | Oblong | Many | Continuous | Orange |
| Nilima | long | 5-10 | Green | Acute | Acute | Conspicuous | Spindle | Many | Continuous | Orange |
| Tiyasha | long | 5-10 | Green | Acute | Acute | Non-Conspicuous | Spindle | Many | Continuous | Orange |
| BG-OP-008 | Extra long | 5-10 | Dark Green | Acute | Acute | Non-Conspicuous | Spindle | Medium | Continuous | Orange |
| BG-OP-002 | Extra long | 5-10 | Glossy Green | Acute | Acute | Non-Conspicuous | Spindle | Few | Continuous | Orange |
| Katahi | Extra long | >10 | Green | Acute | Acute | Conspicuous | Spindle | Many | Discontinuous | Orange |
| Bolder Uchaa | Medium | 5-10 | Glossy Green | Obtuse | Acute | Conspicuous | Oblong | Many | Discontinuous | Orange |
| Kalasona | Long | 5-10 | Light Green | Acute | Acute | Non-Conspicuous | Spindle | Medium | Discontinuous | Orange |
| RS Meghana | Long | 5-10 | Green | Acute | Acute | Non-Conspicuous | Spindle | Medium | Continuous | Orange |
| 2015 Max | Medium | 5-10 | Glossy Green | Obtuse | Acute | Conspicuous | Spindle | Many | Discontinuous | Orange |
| Bolder | Medium | 5-10 | Glossy Green | Obtuse | Acute | Conspicuous | Spindle | Many | Continuous | Orange |
| Diamond | Medium | 5-10 | Green | Obtuse | Acute | Conspicuous | Spindle | Many | Discontinuous | Orange |
| Kajalgouri | Extra long | >10 | Dark Green | Acute | Acute | Conspicuous | Spindle | Many | Continuous | Orange |
| Kalisankar | Long | >10 | Dark Green | Acute | Obtuse | Conspicuous | Spindle | Many | Continuous | Orange |
| Nilisha Gold | Extra long | 5-10 | Glossy Green | Acute | Acute | Non-Conspicuous | Spindle | Many | Continuous | Orange |
| SP Daspur | Long | >10 | Dark Green | Acute | Obtuse | Conspicuous | Spindle | Many | Discontinuous | Orange |
| SP Dhanakali | Extra long | 5-10 | Dark Green | Acute | Acute | Conspicuous | Spindle | Many | Discontinuous | Orange |
| Tiyasha Double | Extra long | 5-10 | Glossy Green | Acute | Acute | Non-Conspicuous | Spindle | Medium | Continuous | Orange |
| Summer special | Long | 5-10 | Glossy Green | Acute | Acute | Conspicuous | Spindle | Many | Continuous | Orange |
| Chaman | Extra long | >10 | Glossy Green | Acute | Obtuse | Conspicuous | Spindle | Many | Continuous | Orange |
| Jonpuri | Extra long | >10 | Dark Green | Acute | Acute | Non-Conspicuous | Spindle | absent | Continuous | Orange |

For peduncle length, 68% of the varieties exhibited lengths ranging between 5 cm and 10 cm, while 32% had peduncles measuring more than 10 cm. Interestingly, no variation was observed for flower Colour, as all varieties uniformly exhibited yellow flowers. However, variation was detected in ovary length, with 18% of the varieties categorized as short (<1.5 cm), 59% as medium (1.6 cm–2.5 cm), and 23% as long (>2.5 cm) (Table 6; Figure 2).

Fruit shape at base was distributed as 77% and 33% between acute and obtuse, respectively. No fruits were found to have a rounded or flattened base. Similarly, the fruit shape at the apex was acute in 86% of the varieties, with the remaining 14% being obtuse, and no instances of a rounded or flattened apex were noted. In longitudinal section, 86% of the fruits were spindle-shaped, while 14% were oblong. Fruit shape was predominantly spindle for the observed genotype indicating the narrowing down of variability with respect to fruit shape owing to preference towards particular shape and selection in similar direction. A significant variation was observed in the presence of fruit tubercles, ranging from absent to many. The majority (68%) had many tubercles, 23% had a medium number, while only 5% fell into each of the categories of absent and few. Regarding tubercle conspicuousness, 64% of the fruits had conspicuous tubercles, while 36% had non-conspicuous ones (Table 6; Figure 3). Fruit ridges were found to be continuous in 68% of the varieties, whereas 32% exhibited discontinuous ridges (Figure 4). Interestingly, no variation was observed in fruit colour at the ripening stage, as all fruits were uniformly orange.

**Figure 3.**
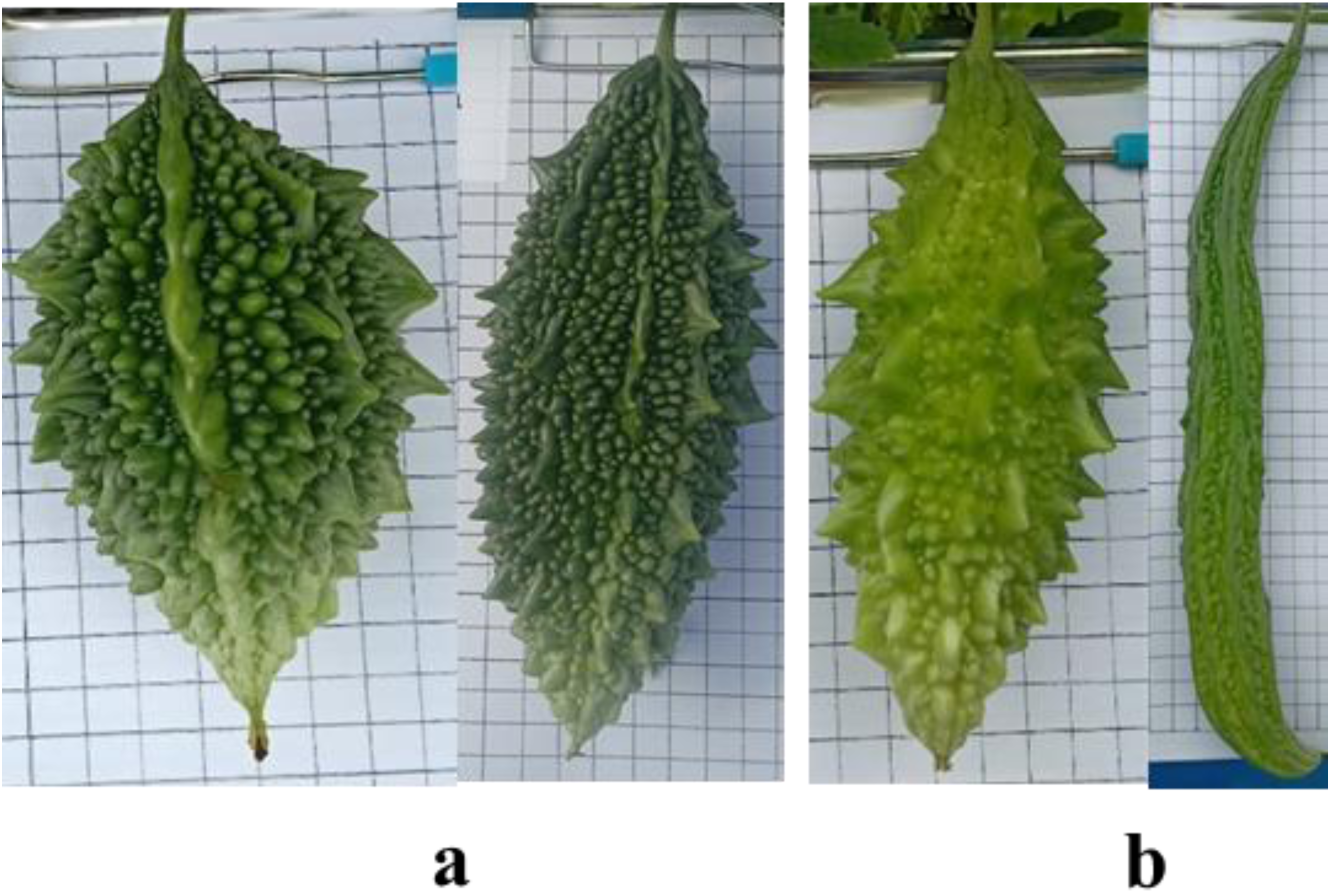
Variation in Fruit Tubercles Prominence: (a) Conspicuous, (b) Non-conspicuous. 34

**Figure 4.**
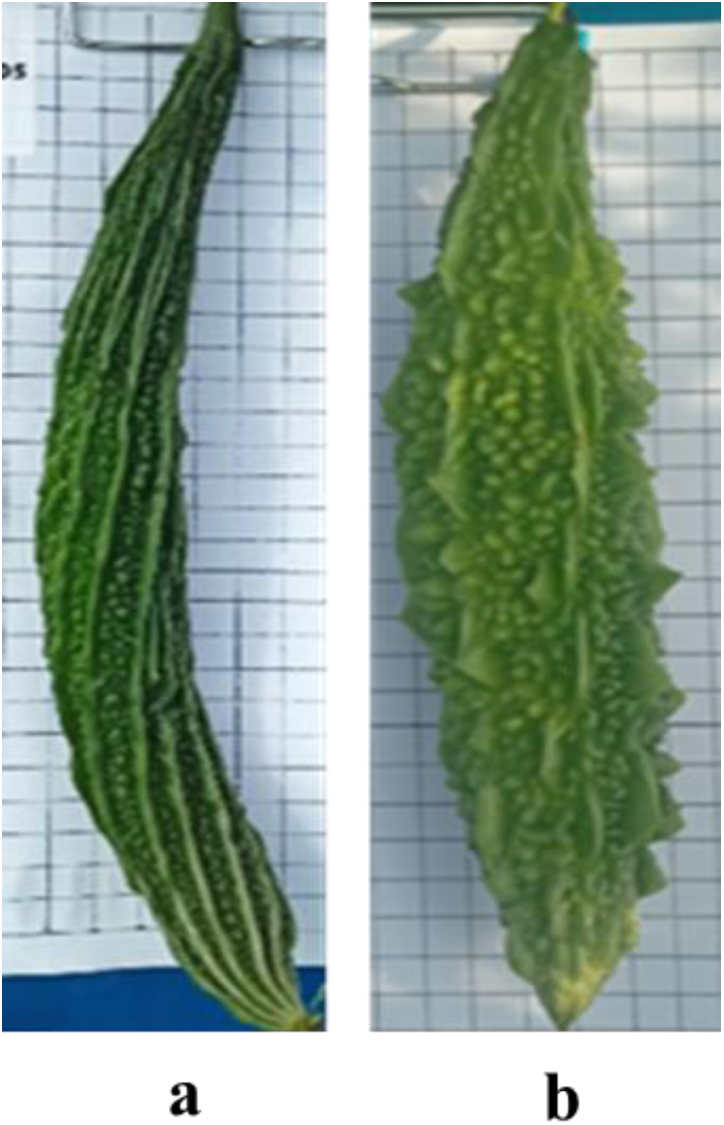
Variation in Fruit Ridge: (a) Continuous, (b) Discontinuous.

### Seed Characteristics

In the present investigation, five traits viz., Number of seeds per fruit, seed length, seed Colour, seed surface and indentation of margin were documented for seed related characteristics as per the DUS guideline. Out of the five, seed surface was found to be monomorphic as all of the collections exhibited a rough surface. Details of trait expressed by different collections (Table 7). The number of seeds per fruit varied from low to medium, with 36% of the samples falling into the low category and 64% classified as medium. Seed length showed considerable variation, with 32% of the seeds being short (<1.4 cm) and the remaining 68% being long (>1.4 cm). Seed Colour was predominantly brown, observed in 77% of the samples, while the remaining 23% were light brown. A notable variation was observed in the indentation of the seed margin, which ranged from small to large. Among the studied samples, 41% had small indentation, 27% had medium, and 32% had large indentation. Notably, no variation was detected for seed surface, as all seeds had a rough texture.

**Table 7:** Characterization of Bitter Gourd Genotypes for Seed Traits According to DUS Guidelines.

| **Characteristics** | **Seediness (no. of seeds/fruit)** | **Seed: length** | **Seed: colour** | **Seed: indentation of margin** |
| --- | --- | --- | --- | --- |
| Kajal | Medium (21-30) | Short | Brown | Large |
| Kuheli | Medium (21-30) | Long | Light Brown | Small |
| Nilima | Less (10-20) | Long | Brown | Large |
| Tiyasha | Less (10-20) | Long | Light Brown | Small |
| BG-OP-008 | Less (10-20) | Short | Brown | Medium |
| BG-OP-002 | Medium (21-30) | Short | Brown | Medium |
| Katahi | Medium (21-30) | Short | Brown | Medium |
| Bolder Uchaa | Less (10-20) | Long | Brown | Small |
| Kalasona | Medium (21-30) | Long | Brown | Large |
| RS Meghana | Medium (21-30) | Long | Brown | Large |
| 2015 Max | Less (10-20) | Short | Brown | Large |
| Bolder | Less (10-20) | Long | Brown | Large |
| Diamond | Less (10-20) | Long | Brown | Medium |
| Kajalgouri | Medium (21-30) | Long | Brown | Small |
| Kalisankar | Medium (21-30) | Short | Brown | Medium |
| Nilisha Gold | Medium (21-30) | Long | Light Brown | Small |
| SP Daspur | Less (10-20) | Long | Brown | Small |
| SP Dhanakali | Medium (21-30) | Long | Light Brown | Small |
| Tiyasha Double | Medium (21-30) | Long | Light Brown | Large |
| Summer special | Medium (21-30) | Short | Brown | Small |
| Chaman | Medium (21-30) | Long | Brown | Medium |
| Jonpuri | Medium (21-30) | Long | Brown | Small |

### Uniformity of varieties

The high degree of cross-pollination in Bitter Gourd can lead to admixtures within the population. This means that different genetic variations can mix freely, creating opportunities for new combinations of traits at the chromosomal level. This natural process offers valuable potential for developing new genetic variations, but it also poses challenges. In this study, out of the 23 genotypes analyzed, 15 genotype displayed consistent phenotypic traits. However, the remaining eight varieties—Summer Special, Tiyasha Double, Nilisha Gold, Kajal Gouri, Kalisankar, Jonpuri, BGOP-Chaman, and 2015 MAX—showed signs of admixture. This finding indicates that these local varieties could benefit from further purification to enhance the consistency of their desirable traits.

## DISCUSSION

Our study revealed significant variability within Bitter Gourd morphological characters, providing a strong foundation for further genetic enhancement of this crop. This observation corresponds with the findings of Behera *et al*. (2008) and Dey *et al*. (2006), who utilized RAPD and ISSR markers to demonstrate genetic distinctness among Bitter Gourd accessions. Among the studied traits, fruit colour, length, tubercles, ridges, seed colour, and indentation of seed margins exhibited the most significant variations, proving highly valuable for differentiating varieties. This finding is consistent with the conclusions of Suma *et al*. (2022) and Mallikarjuna *et al*. (2024), who emphasized the greater reliability of fruit and seed characteristics compared to leaf and flower traits for varietal distinction. A similar study indicated that fruit weight is the primary contributor, followed by fruit yield and fruit length (Rathod *et al*., 2021; Saeed *et al*., 2024). Significant variation has also been reported in key agronomic traits, such as days to flowering, fruit size, and shape, highlighting the abundant genetic diversity within Bitter Gourd germplasm. Similarly, in melon (*Cucumis melo*), fruit characteristics have been identified as key factors for categorizing varieties into distinct groups (Choudhary *et al*., 2015; Dhillon *et al*., 2016; Pandey *et al*., 2019; Alhariri *et al*., 2021). In the present study, no significant variation in vine length was observed, a result consistent with the previous findings (Singh *et al*., 2013). This observation contrasts with the findings of previous studies, which identified vine length as a significant trait contributing to genetic divergence (Jatav *et al*., 2022; Panigrahi *et al*., 2024). The discrepancy may be attributed to differences in sample size and the geographical origin of the accessions analyzed in these studies. Interestingly, four of the 22 collections exhibited characteristics typical of smaller-sized Bitter Gourd varieties. A thorough literature review highlighted the significant contributions of a research group in *Momordica* research from 2005 to 2009, focusing on the collection and characterization of landraces. Their work classified *Momordica* into two botanical varieties: *M. charantia var. charantia* (large-fruited) and *var. muricata* (small-fruited, wild type) (Joseph, 2005; Joseph & Antony, 2008; 2009). Other researchers further validated this classification by demonstrating genetic differences between these two varieties through marker analysis (Behera *et al*., 2008). Recognizing the unique characteristics of the smaller-fruited variety, Suma *et al*. developed a dedicated set of descriptors for *M. charantia var. muricata*, facilitating the identification and characterization of diverse small Bitter Gourd collections (Dey *et al*., 2006). Based on these findings, we tentatively classify the four collections (Bolder, Bolder Uccha, Diamond, and 2015-Max) as belonging to *M. charantia var. muricata*.

In the present study, admixture was observed in seven out of the 22 varieties under investigation, making them unsuitable for commercial production until further purification is conducted Choudhary *et al*. (2015) also emphasized the strict maintenance breeding of reference varieties, including example varieties, to prevent admixture and recommended the use of alternate example varieties for conducting DUS testing in muskmelon. Our findings highlight the significant genetic diversity within Bitter Gourd germplasm, which can be leveraged for future breeding programs aimed at improving agronomic traits. However, the observed admixtures in some varieties suggest the need for further purification efforts to maintain varietal consistency. Future studies should focus on genetic characterization using molecular markers to better understand the genetic basis of these traits and further refine the classification of Bitter Gourd varieties. Additionally, incorporating molecular breeding techniques could help enhance desirable traits such as fruit size, yield, and disease resistance, ultimately benefiting both commercial production and consumer preferences.

## CONCLUSION

The DUS characterization analysis conducted on Bitter Gourd genotypes from East India revealed significant variation, encompassing both *M. charantia var. charantia* and *M. charantia var. muricata*, highlighting the region’s rich genetic diversity. This diverse germplasm represents a valuable resource for breeders, offering opportunities to select inbred lines tailored to specific breeding objectives. The variation observed in traits such as leaf blade margin, immature fruit Colour, fruit length, fruit tubercles, and ridges proved to be the most effective in distinguishing between varieties. These traits hold considerable potential for enhancing cultivar improvement and germplasm conservation efforts, ultimately contributing to increased productivity of Bitter Gourd. However, the current DUS system has limitations, particularly in differentiating between Smooth-type (e.g., Jhalari) and Spiny-type Bitter Gourd. To address this gap, future research should focus on developing more inclusive DUS criteria that better account for the observed morphological diversity. This could involve incorporating additional traits such as the presence or absence of fruit ridges and tubercles. Such modifications would allow for more accurate differentiation between Smooth and Spiny varieties of Bitter Gourd. It provides clear guidelines for identifying and characterizing Bitter Gourd genotypes. Enhancing the effectiveness of DUS testing will contribute to broader goals in cultivar development, market standardization, and germplasm conservation. Future breeding programs could also explore molecular techniques like marker-assisted selection to refine the genetic basis of desirable traits, ultimately leading to the production of high-yielding, disease-resistant, and market-preferred varieties of Bitter Gourd.

## ACKNOWLEDGEMENTS

The author would like to acknowledge Bayer Science and Innovation Private Limited (BSIPL), Bangalore, India for their technical support.

## CONFLICT OF INTEREST

All authors declare no conflict of interest.

## FUNDING

The author(s) received no financial support for the research and publication of this article.

## ETHICS STATEMENT

Not applicable.

## Notes

### Competing Interest Statement

The authors have declared no competing interest.

